# Perinatal exposure to lead results in altered DNA methylation in adult mouse liver and blood: Implications for target versus surrogate tissue use in environmental epigenetics

**DOI:** 10.1101/783209

**Authors:** LK Svoboda, K Neier, R Cavalcante, Z Tsai, TR Jones, S Liu, JM Goodrich, C Lalancette, JA Colacino, MA Sartor, DC Dolinoy

## Abstract

**Background:** DNA methylation is a critical epigenetic mechanism linking early developmental environment to long-term health. In humans, the extent to which toxicant-induced changes in DNA methylation in surrogate tissues, such as blood, mirror those in the target tissues is unclear. The Toxicant Exposures and Responses by Genomic and Epigenomic Regulators of Transcription (TaRGET II) consortium was established by the National Institute of Environmental Health Sciences to address the utility of surrogate tissues as proxies for toxicant-induced epigenetic changes in target tissues.

**Objectives:** The objective of this study was to investigate the effects of perinatal exposure to a human environmentally relevant level (32 ppm in maternal drinking water) of lead (Pb) on liver and blood DNA methylation in adult male and female mice. We hypothesized that developmental Pb exposure would lead to persistent changes in DNA methylation, and that a subset of differentially methylated loci would overlap between liver and blood.

**Methods:** Enhanced reduced-representation bisulfite sequencing was used to assess DNA methylation in 5 month old Pb-exposed and control mice. Sex-stratified modeling of differential methylation by Pb exposure was conducted using an established bioinformatics pipeline.

**Results:** Although Pb exposure ceased at 3 weeks of age, we observed thousands of stably modified, sex-specific differentially methylated regions in the blood and liver of Pb-exposed animals, including 44 genomically imprinted loci. In males, we discovered 5 sites that overlapped between blood and liver, and exhibited changes in DNA methylation in the same direction in both tissues.

**Conclusions:** These data demonstrate that perinatal exposure to Pb induces sex-specific changes in hepatic DNA methylation in adulthood, some of which are also present in blood. Ongoing studies will provide additional exposure-specific insights, and include other epigenetic marks that will enable further refinement of the design and analysis of human studies where target tissues are inaccessible.

## INTRODUCTION

It is increasingly evident that environmental exposures and nutritional perturbations during critical windows of development can influence the long-term risk of disease. One important mechanism by which early-life environment can reprogram disease risk is *via* modification of the epigenome. Normal human development is governed by precise spatio-temporal regulation of gene expression, and this is driven in large part by epigenetic mechanisms. Methylation of DNA at the 5th-position of cytosines (5-methylcytosine, 5mC) occurring in cytosine-phosphate-guanine (CpG) dinucleotides is critical for establishment of tissue-specific gene expression patterns, regulation of imprinted genes, maintenance of genome stability, and silencing of transposable elements (Jones 2012). 5mC undergoes distinct waves of reprogramming during early embryonic development and in primordial germ cells, and is stable and mitotically heritable (Reik et al. 2001). Epigenetic reprogramming during these periods of rapid growth and differentiation can be altered by environmental or nutritional perturbation. 5mC can undergo further oxidation to 5-hydroxymethylcytosine (5hmC) (Lopez et al. 2017). Although 5hmC is an intermediate in the DNA demethylation pathway, recent studies suggest that the genomic distribution, downstream targets, and functional roles of 5hmC are distinct from that of 5mC (Lopez et al. 2017). Environmental exposures may also lead to reprogramming of 5hmC, and these changes are associated with altered transcription and adverse health outcomes (Huang et al. 2019; Kochmanski et al. 2018; Ringh et al. 2019).

Genomic imprinting, which results in mono-allelic, parent-of-origin expression of a subset of genes, is mediated in large part by DNA methylation, and establishment of 5mC at imprinted loci occurs during early embryonic development (Barlow and Bartolomei 2014; Okae et al. 2014). Importantly, we and others have reported that environmental exposures during this critical developmental window lead to altered 5mC and 5hmC at imprinted loci (Bansal et al. 2017; Faulk et al. 2014b; Kochmanski et al. 2018; Nye et al. 2016; Susiarjo et al. 2013) and alterations in imprinted gene expression (Kochmanski et al. 2018; Susiarjo et al. 2013). Abnormal DNA methylation and expression patterns of imprinted genes are associated with several diseases, including growth, metabolic and developmental disorders, cardiovascular diseases, psychiatric illnesses, and cancer (Skaar et al. 2012).

Exposure to heavy metals, including lead (Pb), remains a significant public health concern, particularly in poor urban communities due to various sources of contamination (e.g., leaded paint and pipes, mining, lead-glazed ceramics). Although the neurological effects of early-life Pb exposure have been well-documented, substantially less is known about how Pb exposure impacts the liver and other metabolic organs. Notably, recent human studies suggest that blood Pb levels are associated with increased risk of liver disease and diabetes (Bener et al. 2001; Cave et al. 2010; Zhai et al. 2017). Likewise, maternal blood Pb levels are associated with increased DNA methylation of the imprinted *MEG3* locus in offspring blood, concomitant with rapid gains in body fat (Nye et al. 2016). In animal models, we and others have recently reported that perinatal (Faulk et al. 2013; Faulk et al. 2014a; Leasure et al. 2008; Wu et al. 2016) and post-natal (Sun et al. 2017; Xia et al. 2018a; Xia et al. 2018b) exposure to Pb leads to sex-specific changes in body weight, metabolic function, and gut microbiota. However, although *post-natal* Pb exposure leads to hypermethylation of genes associated with glucose and lipid metabolism in the rat liver (Sun et al. 2017), the effects of perinatal Pb exposure on hepatic DNA methylation, and the implications this may have for long-term disease risk, are unclear.

In human population-based environmental epigenetics studies, easily obtainable sources of DNA (e.g. hair, blood, stool, saliva) are used as surrogates for the tissue(s) targeted by the environmental exposure. It is unknown, however, to what extent exposure-induced epigenetic changes in surrogate tissues reflect those occurring in target tissues. Likewise, whether toxicant-induced epigenomic changes persist in both target and surrogate tissues over time within individuals is unclear. To address these important questions, the National Institute of Environmental Health Sciences (NIEHS) established the Toxicant Exposures and Responses by Genomic and Epigenomic Regulators of Transcription (TaRGET II) consortium (Wang et al. 2018). Using mouse models of human-relevant environmental exposures, the objective of TaRGET II is to assess the conservation of environment-induced epigenomic perturbations between easily obtainable surrogate and disease-relevant, but inaccessible, target tissues. As part of this effort, we used an established mouse model of perinatal Pb exposure to determine the effects of Pb on genome-wide DNA methylation in the liver and blood of 5-month old, adult animals. We hypothesized that perinatal Pb exposure would lead to alterations in DNA methylation in liver and blood that persist into adulthood, and that a subset of these sites would overlap between the two tissues. Additionally, given work from our lab and others demonstrating that Pb exposure is associated with alterations in 5mC at imprinted loci (Faulk et al. 2014b; Nye et al. 2016), we hypothesized that these loci would be targets of Pb exposure in our study.

## METHODS

### Animal Exposure Paradigm

Mice utilized for breeding and exposure were obtained from a colony maintained for over 230 generations with the *A*^*vy*^ allele passed through the male line, resulting in forced heterozygosity on a genetically invariant background with 93% identity to C57BL/6J (Waterland and Jirtle 2003; Weinhouse et al. 2014). Virgin *a/a* females (6–8 wks old) were mated with virgin *a/a* males (7–9 wks old), and randomly assigned to control or Pb exposure via drinking water. For Pb exposure, Pb-acetate was mixed with water to result in a Pb concentration of 32 ppm in drinking water, to model human relevant maternal exposure in the 16-60 μg/dL range (Faulk et al. 2013). Exposure water was made by dissolving Pb (II) acetate trihydrate (Sigma-Aldrich) in a single batch of distilled water, and Pb concentrations were verified using inductively coupled plasma mass spectrometry with a limit of detection of 1.0 µg/L (ICPMS; NSF International). Animals were maintained on a phytoestrogen-free modified AIN-93G diet (TD.95092, 7% Corn Oil Diet, Harlan Teklad). The *a/a* dams in the Pb group were exposed to Pb-supplemented drinking water for two weeks prior to mating *a/a* males. Exposure was continued during gestation and lactation. After weaning, the resulting pups were weighed and switched to Pb-free drinking water **(Figure 1).** A subset of pups from each exposure group, representing approximately 1–2 male and 1–2 female offspring per litter, was maintained to 5 months of age (n=6 males and 6 females per treatment). All animals had access to food and drinking water *ad libitum* throughout the experiment while housed in polycarbonate-free cages. The study protocol was approved by the University of Michigan Institutional Animal Care and Use Committee (IACUC).

**Figure 1:**
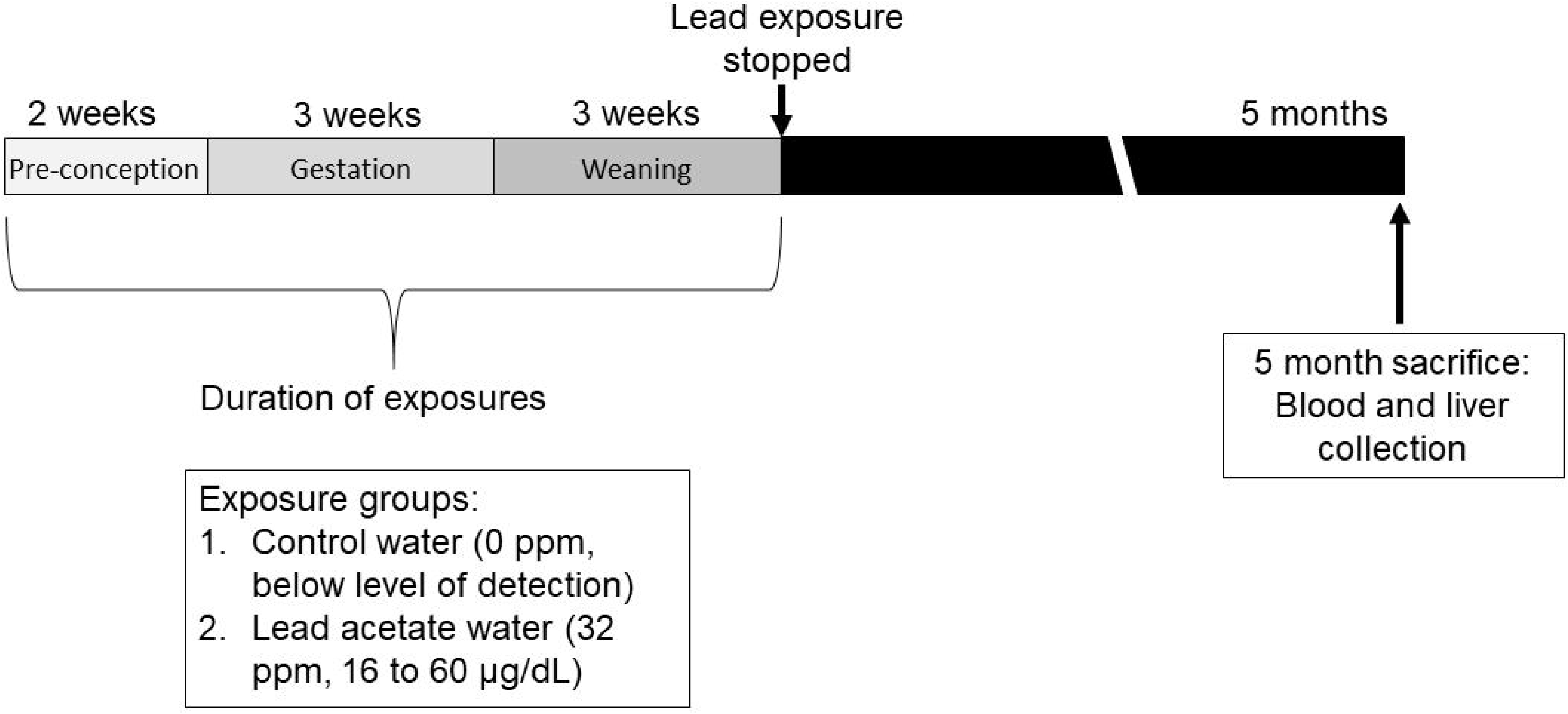
Mouse treatment paradigm showing timing of Pb exposure and animal sacrifice/tissue collection. Dams were exposed to normal chow and either control or Pb-acetate water, starting two weeks prior to mating, through gestation, until offspring were weaned on postnatal day 21. At this point, Pb exposure ceased and offspring were administered control water and chow. At 5 months of age, matched liver and blood tissues were collected for DNA methylation analysis. Pb doses and average blood lead levels are shown.

### Body Composition and Tissue Collection

Body weights were measured for each mouse on a weekly basis (Mettler Toledo). Total body fat and lean mass were measured via EchoMRI (Houston, TX USA). Body fat percentage and lean mass percentage were calculated as a percent of total body weight. Upon euthanasia at 5 months of age, mesenteric fat, blood, and liver samples were collected in accordance with protocols established by the TaRGET II Consortium (**Figure 1** and (Wang et al. 2018)). Briefly, mice were fasted for 6 hours prior to euthanasia. Euthanasia was carried out via CO_2_ asphyxiation and bilateral pneumothorax. Blood was collected by cardiac puncture, followed by whole body perfusion with cell culture grade 0.9% saline solution (Sigma Life Sciences). Next, liver and mesenteric fat were collected and weighed. Mesenteric fat was collected by dissecting white adipose tissue away from the stomach, spleen, pancreas, and intestines. Relative liver and mesenteric fat weights were expressed as a percent of total body weight. Liver was separated into individual lobes, and the left lobe was used for the analyses in this study. Liver and fat samples were immediately snap frozen in liquid nitrogen and stored at −80°C. Blood was collected with EDTA anticoagulant, centrifuged, and plasma was removed. Red blood cells were lysed using Erythrocyte Lysis Solution (Qiagen, 79217). The remaining white blood cell fraction was washed with PBS, pelleted by centrifugation, and resuspended in Buffer RLT (Qiagen, cat # 1053393). Crude cell lysate was stored at −80°C until DNA extraction.

### DNA Extraction and Enhanced Reduced Representation Bisulfite Sequencing

The left lobe of the liver was cryo-pulverized and suspended in Buffer RLT (Qiagen, cat # 1053393). Blood and liver DNA extraction were performed using the AllPrep DNA/RNA/miRNA Universal Kit (Qiagen #80224). Enhanced reduced representation bisulfite (ERRBS) was performed at the University of Michigan Epigenomics Core as described previously (Akalin et al. 2012; Garrett-Bakelman et al. 2015). Briefly, 50 ng of genomic DNA was digested using MspI, a restriction enzyme that preferentially cuts CG-rich sites. The digested DNA was then purified using phenol:chloroform extraction and ethanol precipitation in the presence of glycogen, before blunt-ending and phosphorylation. A single adenine nucleotide was next added to the 3′ end of the fragments in preparation for the ligation of the adapter duplex with a thymine overhang. The ligated fragments were cleaned, then processed for size selection on agarose gel. Selected fragments were treated with sodium bisulfite to convert unmethylated cytosines to uracils, which are then replaced with thymines during PCR amplification. Both methylated and hydroxymethylated CpGs are protected from bisulfite conversion. Therefore, ERRBS detects a combination of 5mC and 5hmC. Final libraries were next cleaned up with AMPure XP beads (Product #A63880; Beckman Coulter), quantified using the Agilent TapeStation genomic DNA kit (Catalog #G2991AA; Agilent) and Qubit High Sensitivity dsDNA (Catalog #Q32850; Invitrogen). Single-end, 50 bp reads were obtained for each library by sequencing on the HiSeq 4000 system (Illumina). ERRBS samples were multiplexed, with three samples per sequencing lane.

### Bioinformatics pipeline, quality control, and differential methylation analysis

Snakemake (Koster and Rahmann 2012) was used to manage the bioinformatics workflow, and Anaconda (Anaconda Software Distribution) was used to manage software dependencies and create reproducible software environments. For quality control, FastQC (v0.11.3) was employed to assess the overall quality of each sequenced sample and identify specific reads and regions for trimming. TrimGalore (v0.4.5) was used to trim low-quality bases (quality score lower than 20), adapter sequences (required overlap of 6bp), and end-repair bases from the 3’ end of reads. Reads shorter than 20bp after trimming were discarded. For alignment and methylation calling we used Bismark (Krueger and Andrews 2011) (v0.19.0), an integrated alignment and methylation call program that performs unbiased alignment (by converting residual cytosines to thymines prior to alignment in both reads and reference). Within Bismark, reads were aligned to the reference genome (mm10) using Bowtie2 (Langmead and Salzberg 2012) (v2.3.4) with default parameter settings (multi-seed length of 20bp with 0 mismatches). Methylation calls were reported for all nucleotides with a read depth of at least 5. We used the methylSig R package (Park et al. 2014) (v0.5.0) to identify differentially methylated CpG positions and/or regions. Differential methylation for each comparison was tested using methylSigDSS(), which tests for differential methylation under general experimental design using a beta-binomial approach with the ‘arcsine’ link function (Park and Wu 2016). To control for batch effects, run was included as a covariate in the model. After obtaining p-values, we adjusted for multiple testing using the FDR approach. The results of the differential methylation tests were then annotated using the annotatr R Bioconductor package (Cavalcante and Sartor 2017) (v1.5.9). The annotate_regions function was used to generate genomic annotations, which include CpG annotations [CpG islands (CGI), shores, shelves, open sea (InterCGI)], genic annotations (exon, intron, promoter, 5’UTR, 3’UTR), and gene IDs. CpGs with read coverage > 1000 were removed because they were likely the result of PCR amplification, and CpGs with read coverage < 10 were removed due to decreased power to detect differential methylation. Opposite strand CpGs at the same position were combined via destranding. We performed differential methylation testing on individual CpG sites (DMCs), requiring sufficient sequencing coverage for a minimum of 4 samples from the Pb group, and 4 samples from the control group for a site to be tested. Differentially methylated regions (DMRs) were identified in 50 bp tiles using the same requirements. For both DMCs and DMRs, sites with FDR <0.05 and an absolute difference in methylation of > 10% were considered significant.

### Pathway analysis of DMRs

Poly-Enrich (Lee C.T. 2019) was used to assess the biological pathways enriched among the DMRs. Separate pathway analyses were conducted for each sex, tissue, and direction of differential methylation, and were restricted to proximal promoter regions (within1 kb of transcription start sites). The following Gene Ontology gene sets used for this analysis: Biological Process, Cellular Component, and Molecular Function. Pathways with FDR <0.05 were considered statistically significant.

### Analysis of overlap in differentially methylated sites between blood and liver

Annotated lists of DMCs and DMRs for males and females were compared between blood and liver to identify chromosomal locations that directly overlapped between the two tissues. Overlapping sites with changes in methylation in the same direction in both tissues were considered biologically relevant. In a further analysis, the full lists of DMR-associated genes in blood were compared to those in liver to identify a list of genes in common between the two tissues. Separate lists were generated for males and females. These gene lists were analyzed using DAVID version 6.8, Laboratory of Human Retrovirology and Immunoinformatics (LHRI), (https://david.ncifcrf.gov) to identify pathways that were significantly enriched (FDR <0.05).

## RESULTS

### Litter parameters and phenotype

Perinatal lead exposure did not significantly alter the offspring sex ratio, litter size, or mortality rates (p = 1.0, 0.9, and 0.2, respectively, Fisher’s Exact Test) (**Table 1**). We have previously reported that perinatal Pb exposure leads to sex-specific changes in food intake, body weight, and body fat later in life (n=12-18 mice per sex) (Faulk et al. 2014a). However, given the smaller sample size in this study (n=6 mice per sex), it is not surprising that we observed no significant differences in body weight, total body fat, or body fat percentages in either males or females (**Supplemental Figure 1 A-F**). Nevertheless, consistent with our previous observations, we observed a trend toward increased total and relative mesenteric fat weights in males at 5 months, although this did not reach statistical significance (p=0.06, **Supplemental Figure 1 G-J**). Total and relative liver weights did not differ significantly between exposed and control groups in either males or females (**Supplemental Figure 1 K-N**).

**Table 1:** Litter Parameters.

| Exposure Group | # Litters | # Pups Born | # Pups Died | Mean Pups per Litter (± SD)<sup>a</sup> | Pup Mortality Rate (%)<sup>b</sup> | % Females<sup>a</sup> |
| --- | --- | --- | --- | --- | --- | --- |
| Control | 6 | 53 | 3 | 7.57 ± 3.36 | 5.7 | 46% |
| Pb | 7 | 52 | 7 | 7.43 ± 1.51 | 13.5 | 47% |

Note: Statistical analyses (Pb vs. Control) were conducted using Fisher's Exact Test.
<sup>a</sup>p = ≥ 0.9
<sup>b</sup>p = 0.2

### Epigenome-wide differential DNA methylation with Pb exposure

To assess the effects of perinatal Pb exposure on DNA methylation across target and surrogate DNA, ERRBS was conducted on liver and blood samples from 5-month-old Pb and control-exposed offspring (n=6 males and 6 females per exposure group). Tables 2 and 3 illustrate the total number of regions tested and the number of differentially methylated cytosines (DMCs) and DMRs identified in each exposure group. Although Pb exposure ceased at 3 weeks of age, this analysis revealed thousands of sex-specific DMCs and DMRs in the adult liver and blood (**Tables 2-3**). The relative proportion of hypo- and hyper-methylated regions/cytosines was similar for each condition (**Tables 2-3**). Although observed methylation changes were between 10-20% at most sites with Pb exposure, numerous sites exhibited changes in methylation that were much greater, up to 70% in females and 74% in males **(Figure 2 A-D and Supplemental Figure 2)**. We annotated the DMCs and DMRs to the mouse mm10 genome to determine the genomic locations of DMCs and DMRs relative to all sites tested, stratified by sex, tissue, and direction of differential methylation. Compared to all regions tested, DMCs/DMRs were significantly depleted from CpG islands, promoters, 5’UTRs, and exons, and were significantly enriched for regions greater than 4000 bp from a CpG island (interCGI) and intergenic regions (chi squared test, p<.05) (**Supplemental Figure 3 A-H**). This pattern was similar across sex and tissue (**Supplemental Figure 3 A-H**).

**Table 2:** Differentially methylated cytosines (DMCs) in female and male blood and liver identified using ERRBS.

| Condition | Total | # Hypermethylated<br>(% Total) | # Hypomethylated<br>(% Total) | Total Tested |
| --- | --- | --- | --- | --- |
| Female Blood | 1,628 | 747 (45.9) | 881 (54.1) | 1,356,386 |
| Female Liver | 2,404 | 1,350 (56.2) | 1,054 (43.8) | 1,228,997 |
| Male Blood | 2,175 | 1,076 (49.5) | 1,099 (50.5) | 1,362,718 |
| Male Liver | 3,808 | 1,598 (42.0) | 2,210 (58.0) | 1,173,995 |

Note: Methylation changes are expressed as lead vs. control.

**Table 3:** Differentially methylated regions (DMRs) in female and male blood and liver identified using ERRBS

| Condition | Total | # Hypermethylated<br>(% Total) | # Hypomethylated<br>(% Total) | Total Tested |
| --- | --- | --- | --- | --- |
| Female Blood | 1,035 | 551 (53.2) | 484 (46.8) | 646,876 |
| Female Liver | 1,740 | 792 (45.5) | 948 (54.5) | 597,077 |
| Male Blood | 1,404 | 724 (51.6) | 680 (48.4) | 650,782 |
| Male Liver | 3,342 | 1,874 (56.1) | 1,468 (43.9) | 563,681 |

Note: Methylation changes are expressed as lead vs. control.

**Figure 2:**
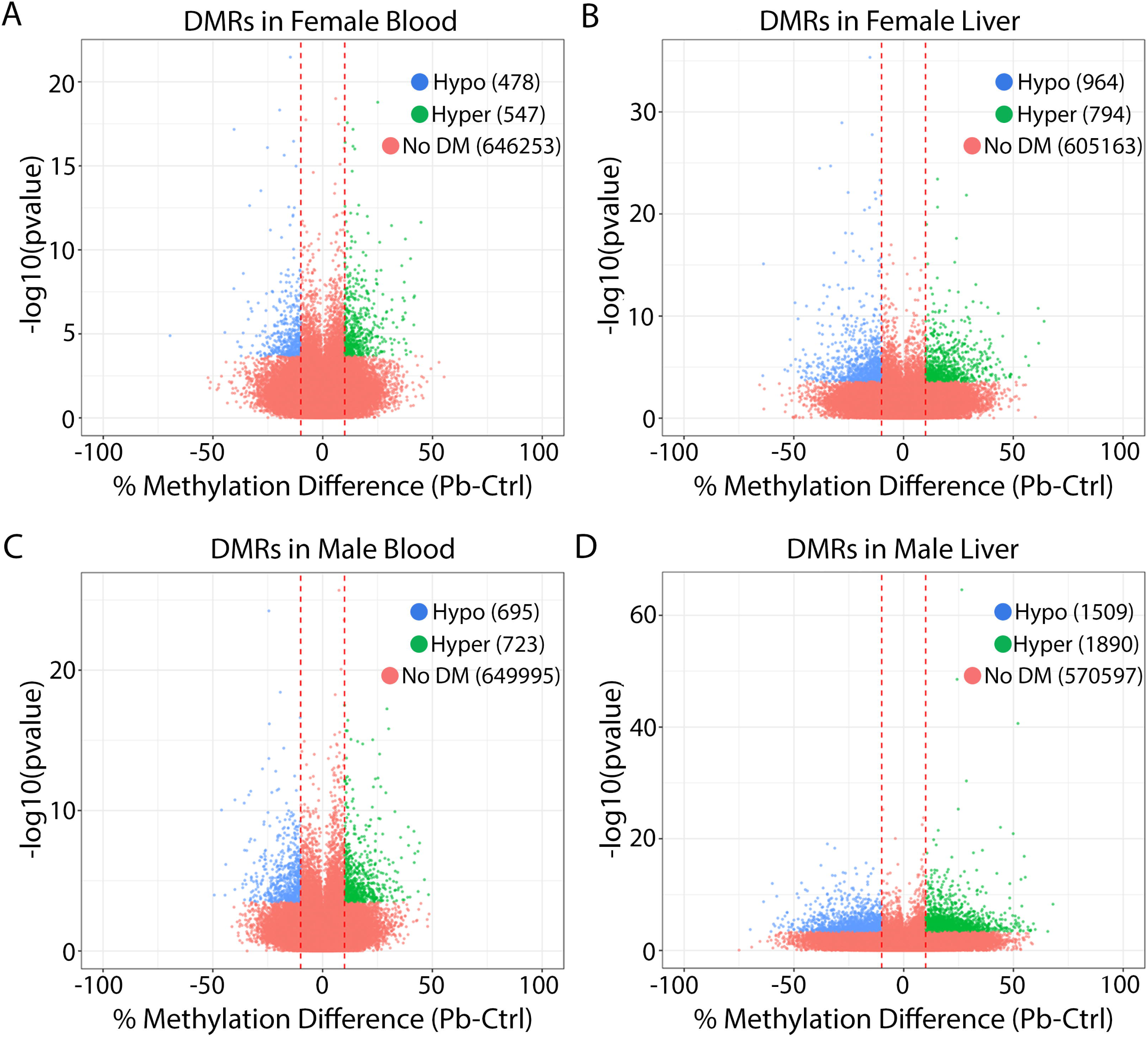
Volcano plots showing differentially methylated regions (DMRs) for lead vs. control in female blood (A), female liver (B), male blood (C), and male liver (D). Red: regions significantly hypermethylated with Pb exposure. Green: regions significantly hypomethylated with Pb exposure.

### Pathway analysis of DMRs

Having established that *early life* Pb exposure leads to changes in DNA methylation in adulthood, we next determined the cellular pathways enriched among the DMRs (**Tables S1-S4**). We stratified our analyses by sex, tissue, and direction of differential methylation, and focused on proximal promoter regions (within1 kb of transcription start sites). Pathways with a FDR<0.05 were considered significant. This analysis revealed that the pathways enriched among hypo-methylated sites were distinct from those enriched among hyper-methylated sites. In addition, the enriched pathways differed across sex and tissue. In spite of these differences, overall, we observed that Pb altered methylation of CpGs in key organ developmental pathways including mammary gland, bone, nervous system, and kidney development. Interestingly, we observed enrichment of pathways involved in regulation of neurotransmitter levels and activity in female and male liver.

### Perinatal Pb exposure leads to changes in methylation at imprinted loci

Given previous findings from our lab and others that developmental exposure to BPA and Pb led to altered CpG methylation at imprinted loci (Faulk et al. 2014b; Kochmanski et al. 2018; Nye et al. 2016), we investigated whether perinatal Pb exposure disrupted methylation at imprinted loci in liver and blood, and whether these changes overlapped between the two tissues. To do this, we compared the lists of Pb-induced DMCs/DMRs for each condition to a database of mouse imprinted genes (Williamson CM 2013). Of 139 imprinted genes interrogated, 112 were covered in this study using ERRBS (see Supplemental Excel file). We observed Pb-induced DMRs/DMCs at 44 imprinted genes combined across tissue and sex. Specifically, we identified 3 in female blood, 18 in female liver, 22 in male blood, and 21 in male liver. See **Tables 4-5** for DMCs mapping to imprinted genes in females and males, respectively, and **Supplemental Tables 5-8** for DMRs. In liver samples, we discovered 5 imprinted genes with DMCs in both males and females, including *Commd1, Gnas, Nespas, Pde10a*, and *Pde4d* (**Tables 4-5**). Methylation at CpGs in *Gnas/Nespas* was in the same direction in both males and females (**Figure 3A**). In blood, CpGs in *Meg3* were hypermethylated in both males and females (**Figure 3B**). We next compared DMC-associated imprinted genes between blood and liver within each sex. In females, no imprinted genes overlapped between blood and liver. In males, we observed DMCs at *Begain, Cdkn1c, Commd1, Peg12, Rasgrf1*, and *Snrpn* in both blood and liver, and found that CpGs in *Begain, Peg12, Rasgrf1*, and *Snrpn* exhibited changes in methylation in the same direction in both blood and liver (**Table 5 and Figure 3D-G**). Collectively, these data suggest that Pb exposure alters methylation at imprinted loci in a tissue and sex-dependent manner.

**Table 4:**
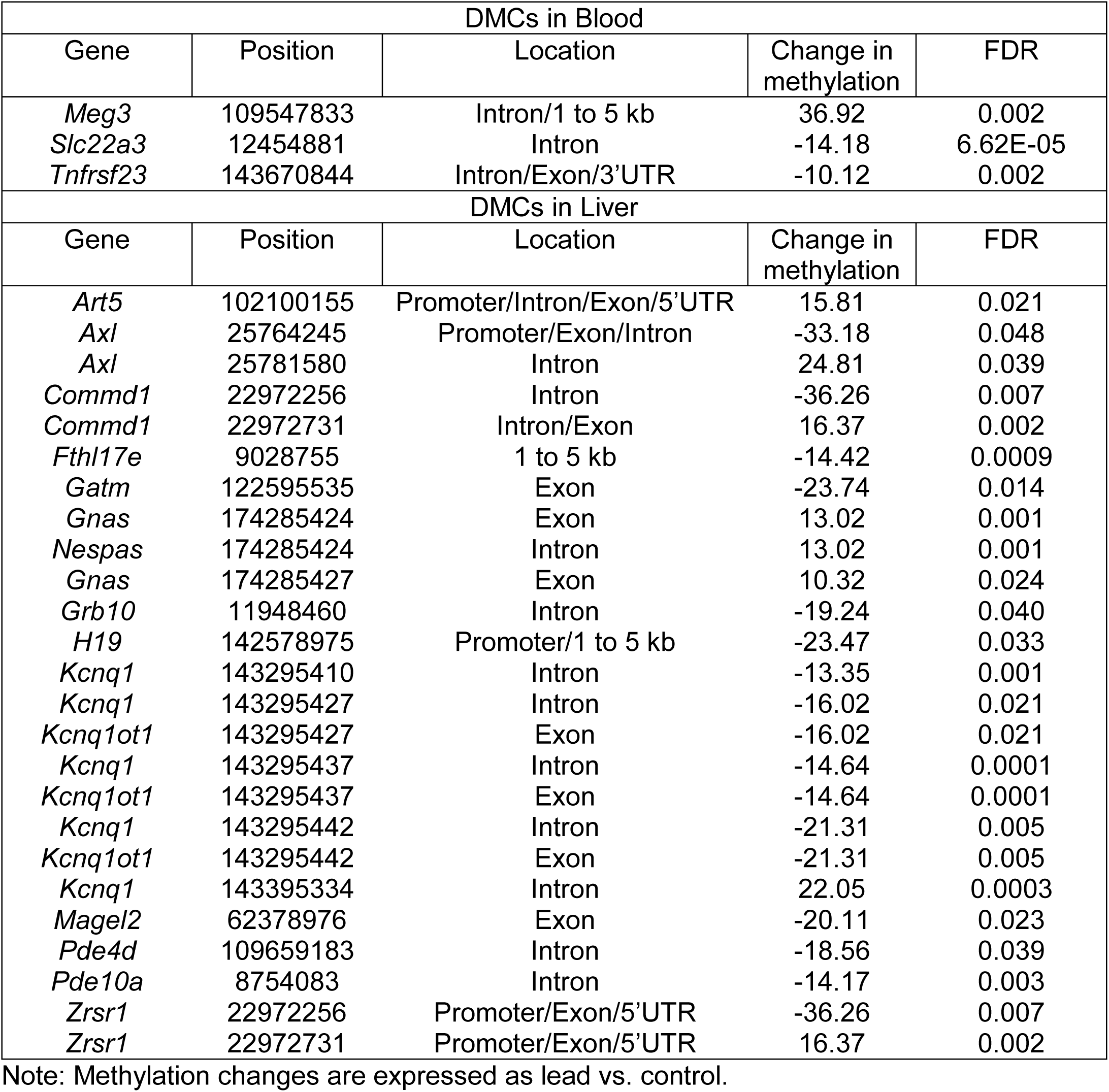
Differentially methylated cytosines (DMCs) identified at imprinted loci in female blood and liver.

**Table 5:**
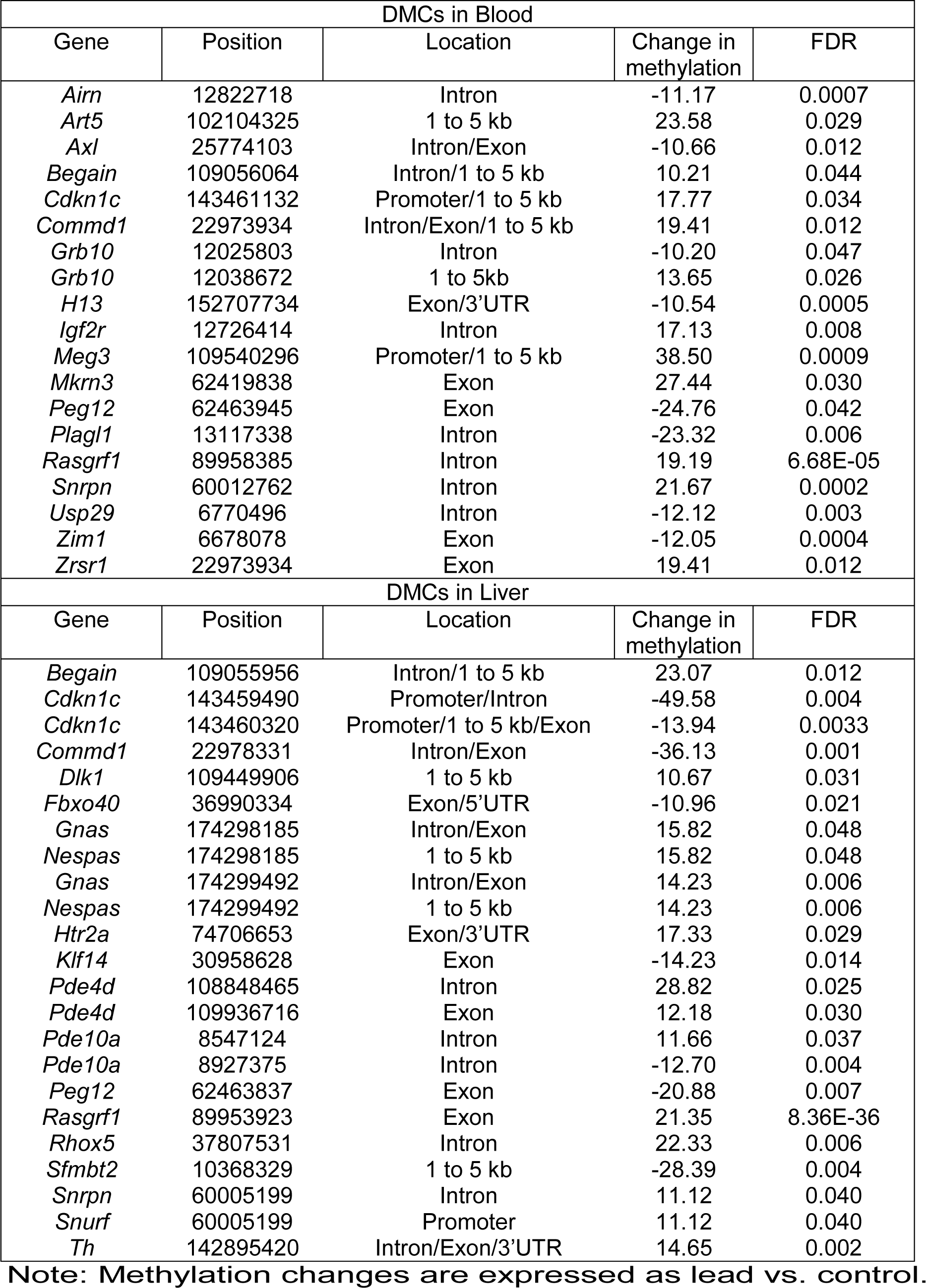
Differentially methylated cytosines (DMCs) identified at imprinted loci in male blood and liver.

**Figure 3:**
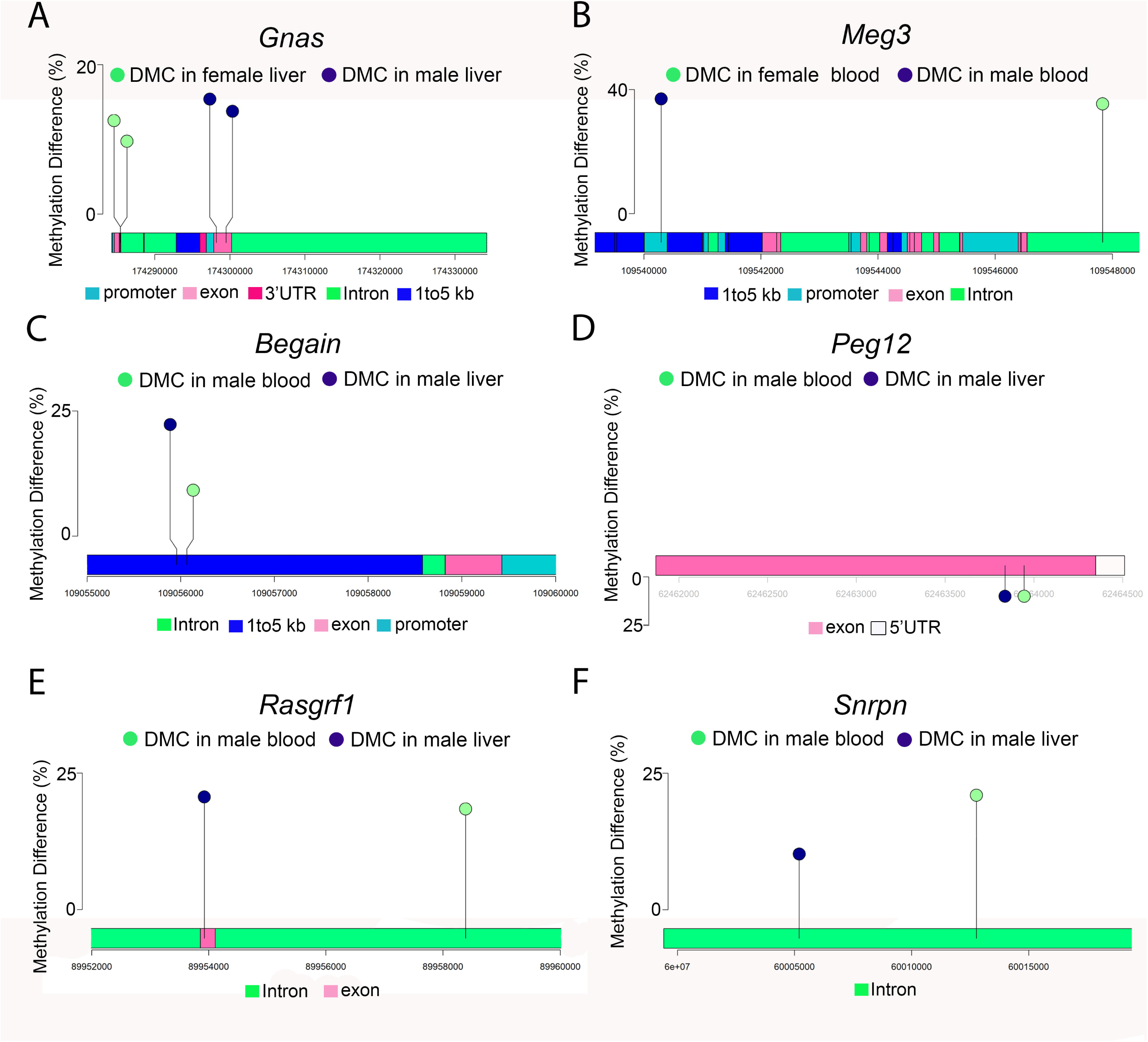
Lollipop diagrams showing differentially methylated cytosines (DMCs) with Pb exposure in: (A) *Gnas* in male and female liver, (B) *Meg3* in male and female blood, (C) *Begain* in male blood and liver, (D) *Peg12* in male blood and liver, (E) *Rasgrf1* in male blood and liver, and (F) *Snrpn* in male blood and liver.

### Overlap of DMRs and DMCs between liver and blood

An important objective of the TaRGET II Consortium is to identify signatures of environmental exposures in surrogate tissues that mirror those in target tissues. To this end, we first compared the chromosomal locations for DMCs and DMRs between liver and blood to determine whether there was overlap in differentially methylated sites between the two tissues. In females, 4 DMCs and 3 DMRs overlapped between blood and liver; however, the Pb-induced change in methylation at each of the overlapping sites was in the opposite direction for each tissue (**Table 6**). In males, 4 DMCs and 5 DMRs each overlapped between blood and liver (**Table 7**). In contrast with females, several sites in males exhibited changes in methylation that were in the same direction in both blood and liver. Among DMCs, these sites mapped to *Grifin, Elmsan1, Amn*, and *Tbc1d30*. Among DMRs, overlapping sites mapped to *Grifin* and *Plekhg3*. An additional region was identified that did not map to a gene (**Table 7**). Next, we evaluated the overlap in DMC-associated genes between blood and liver in males and females. In males and females, we identified 325 genes in males and 188 DMC-associated genes in females that overlapped between blood and liver (**Figure 4 and Supplemental Tables 9-10**). Gene ontology analysis of the overlapping genes revealed enrichment for cell adhesion, protein binding, synapse, and neuronal process pathways (**Figure 4**). Overlapping genes in females were enriched for cell junction, membrane, and synapse pathways. We next compared the lists of DMC-associated genes with blood-liver overlap between males and females. A small number of genes (35 genes) were common between tissues and sexes (**Supplemental Table 11**). Together, these data suggest that there exist sex-specific changes in DNA methylation with perinatal Pb exposure that are consistent across blood and liver in adulthood, with few specific CpG sites overlapping directly between tissues.

**Table 6:** Overlap between differentially methylated cytosines/regions (DMCs/DMRs) in female blood and liver.

| Gene | Coordinate | Change in blood | FDR blood | Change in liver | FDR liver |
| --- | --- | --- | --- | --- | --- |
| *Ccin* | 43985009 | -11.08 | 0.00935 | 28.08 | 0.011 |
| *Lsm14b* | 180023832 | -19.35 | 0.00035 | 16.92 | 0.017 |
| *Cntnap2* | 46349288 | -17.51 | 0.00091 | 39.58 | 0.024 |
| *Fam122c* | 53314658 | -18.76 | 0.00012 | 18.05 | 0.006 |
| Gene | Coordinate | Change in blood | FDR blood | Change in liver | FDR Liver |
| *Glo1* | 30597451 | 11.86 | 0.00092 | -13.38 | 0.045 |
| *Lsm14b* | 180023801 | -19.35 | 0.00026 | 16.92 | 0.012 |
| *Cntnap2* | 46349251 | -17.51 | 0.00070 | 39.58 | 0.018 |

Note: Methylation changes are expressed as lead vs. control.

**Table 7:** Overlap between differentially methylated cytosines/regions (DMCs/DMRs) in male blood and liver.

| DMCs |  |  |  |  |  |  |
| --- | --- | --- | --- | --- | --- | --- |
| Gene | Start | End | Change in blood | FDR blood | Change in liver | FDR liver |
| *Grifin* | 140567250 | 140567250 | -12.83 | 0.025 | -25.08 | 0.03932 |
| *Elmsan1* | 84151896 | 84151896 | 15.08 | 0.0005 | 11.74 | 0.00020 |
| *Amn* | 111270178 | 111270178 | 19.13 | 0.033 | 11.97 | 0.00060 |
| *Tbc1d30* | 121294168 | 121294168 | 22.54 | 0.047 | 27.76 | 0.01649 |
| DMRs |  |  |  |  |  |  |
| Gene | Start | End | Change in blood | FDR blood | Change in liver | FDR liver |
| *Grifin* | 140567201 | 140567250 | -12.83 | 0.020 | -25.08 | 0.026 |
| *Mob2* | 142016401 | 142016450 | -23.64 | 0.038 | 21.0 | 0.021 |
| *Dnm3* | 162453101 | 162453150 | 12.63 | 0.001 | -23.95 | 0.049 |
| *Plekhg3* | 76535801 | 76535850 | 17.04 | 0.017 | 17.14 | 0.047 |
| *N/A* | 114931401 | 114931450 | 21.20 | 0.022 | 13.43 | 0.039 |

Note: Methylation changes are expressed as lead vs. control.

**Figure 4:**
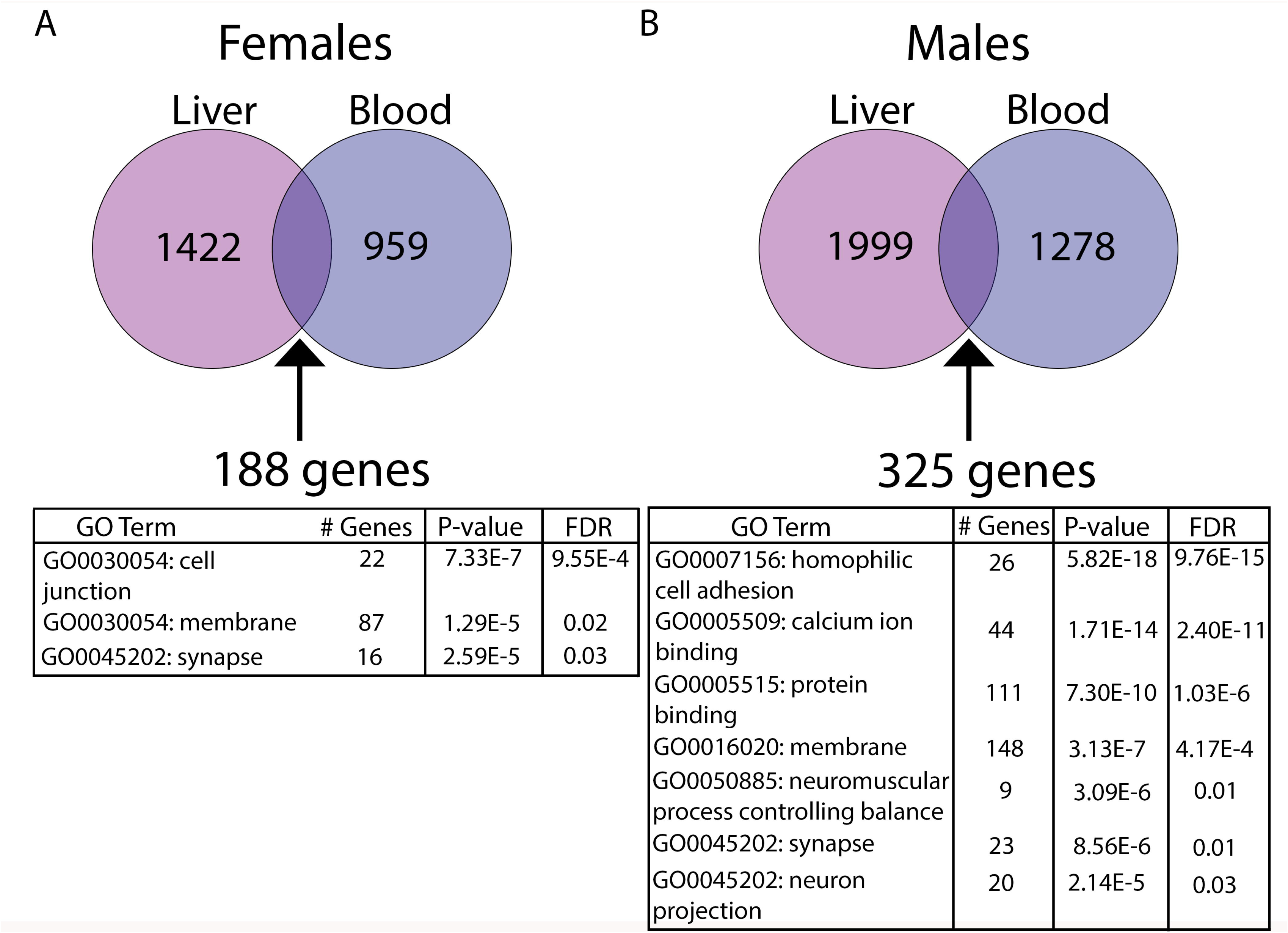
Analysis of overlapping DMC-associated genes in females (A) and males (B). Numbers in circles indicate the total number of DMC-associated genes in each tissue. In the tables, # Genes indicates the number of genes in the specified GO pathway that appear in the experimental dataset. Pathway analysis was conducted using DAVID, version 6.8 (Laboratory of Human Retrovirology and Immunoinformatics (LHRI)).

## DISCUSSION

In this work, we have investigated the effects of perinatal Pb exposure on DNA methylation in the liver and blood of adult mice. The findings from this study are novel for several reasons. First, although increasing evidence links Pb exposure to adverse metabolic outcomes (Niessen et al. 2018), the mechanistic basis for these associations is unclear. Indeed, little is known about the effects of Pb exposure on the liver. To our knowledge, this is the first report demonstrating that Pb exposure during pregnancy and lactation leads to sex-specific alterations in hepatic DNA methylation in adulthood. Second, we discovered that perinatal Pb exposure leads to changes in methylation at several imprinted loci that are also reprogrammed by perinatal BPA exposure (Kochmanski et al. 2018). Finally, we mapped DNA methylation genome-wide in both blood and liver of the same mice after perinatal Pb exposure, and although few differentially methylated sites directly overlapped between blood and liver, we discovered many DMC-associated genes that overlap between tissues in both males and females. This finding has important implications for the design and interpretation of human population-based epigenomics studies.

### Genome-wide changes in DNA methylation with Pb exposure

Although Pb exposure ceased at 3 weeks of age, we discovered thousands of DMCs/DMRs in both male and female mice at 5 months of age. In accordance with previous findings from our lab and others (Faulk et al. 2013; Faulk et al. 2014a; Sen et al. 2015b), the effects of Pb on DNA methylation were highly sex-specific. Indeed, when comparing males and females, pathway analyses revealed enrichment of distinct biological pathways. Likewise, the DMRs/DMCs were mostly tissue type-specific, and the specific sites that directly overlapped between liver and blood differed by sex. Among DMC-associated genes that overlapped between blood and liver, only a small percentage overlapped between males and females. These data suggest that sex will be a critical factor in the design, analysis, and interpretation of human- and animal-based environmental epigenetics studies. Although we investigated the effects of perinatal Pb exposure on DNA methylation in liver and blood, pathway analysis of DMRs in promoter regions revealed enrichment for pathways involved in neurological development and function in male and female liver. Moreover, among overlapping genes between blood and liver, gene pathways associated with synapse function were significantly enriched in both males and females. The nervous system is a well-known target of Pb, and although we did not directly measure nervous system tissues in this study, it is nonetheless striking that neurological pathways were enriched in genes annotated to DMCs/DMRs in both liver and blood. Analysis of other tissues, including brain, from Pb-exposed mice is currently underway, and will provide important insight into whether there is a common subset of tissue-independent, environmentally labile genes or CpG sites.

Although Pb exposure affected DNA methylation at thousands of loci, the mechanism underlying these changes is currently unknown. DNA methyltransferases (DNMTs) methylate the 5-position of cytosine bases in DNA using S-adenosylmethionine (SAM), which is generated via one-carbon metabolism (Ducker and Rabinowitz 2017). Pb may disrupt DNA methylation via alteration in DNMT expression/activity or through perturbation of SAM levels. Consistent with the first hypothesis, Pb exposure *in vitro* and *in vivo* leads to inhibition and altered expression of DNMTs (Sanchez et al. 2017). In support of the second possibility, administration of exogenous SAM mitigates the deleterious effects of Pb exposure, suggesting that Pb may deplete endogenous levels of SAM (Cao et al. 2008; Paredes et al. 1986). As ERRBS cannot discriminate between 5mC and 5hmC, a subset of changes in methylation observed in this study may actually be 5hmC, and altered 5hmC has indeed been reported with Pb exposure (Sen et al. 2015a). 5hmC is generated by the activity of TET methylcytosine dioxygenases (TETs), using the cofactor alpha-ketoglutarate (α-KG), which is generated by the TCA cycle (Wong et al. 2017). Interestingly, Pb exposure has been shown to reduce activity of TCA cycle enzymes, including isocitrate dehydrogenase, which catalyzes the formation of α-KG (Basha et al. 2012; Seddik et al. 2011). Finally, it is possible that developmental Pb exposure leads to shifts in the cellular composition of blood and liver (Hung et al. 2019; Trevino and Katz 2018), and that alterations in DNA methylation are a reflection of these population shifts. The mechanisms underlying the observed Pb-induced changes in DNA methylation are currently under investigation.

### Imprinted Genes

Genomic imprinting is critical for normal fetal development, and epigenetic programming of imprinted gene expression is established very early in development (Dolinoy et al. 2007). For these reasons, this process may be highly susceptible to perturbation by environmental exposures. Notably, we discovered that Pb exposure during gestation and lactation leads to changes in DNA methylation at 44 imprinted genes. This finding is consistent with accumulating evidence from our lab and others demonstrating that early life environmental exposures disrupt the epigenetic programming of imprinted loci (Bansal et al. 2017; Kochmanski et al. 2018; Nye et al. 2016; Susiarjo et al. 2013). Although most of the effects were sex-specific, we observed DMCs at *Commd1, Gnas, Nespas, Pde10a*, and *Pde4d* in both males and females, and methylation changes at CpGs in *Gnas, Nespas, and Pde10a* were in the same direction in both sexes. The *Gnas* locus encodes for three distinct protein products, and disruptions in the regulation of this locus have been implicated in a number of disorders, including early-onset obesity (Hanna et al. 2018). We recently reported that developmental BPA exposure leads to changes in DNA methylation and hydroxymethylation at the *Gnas* locus, concomitant with changes in *Gnas* expression (Kochmanski et al. 2018). Thus, *Gnas* may be an important target of multiple environmental exposures, in both males and females. When comparing male liver and blood, we observed DMCs at *Begain, Cdkn1c, Commd1, Peg12, Rasgrf1*, and *Snrpn* in both tissues, and found that CpGs in *Begain, Peg12, Rasgrf1*, and *Snrpn* exhibited changes in methylation in the same direction in both blood and liver. *Snrpn* encodes for an RNA binding protein that is critical for neurological function, and disruption of *Snrpn* expression is implicated in Prader-Willi syndrome, a neuroendocrine disorder associated with intellectual disability and obesity (Reed and Leff 1994). Work from our lab and others suggests that environmental exposures are associated with changes in methylation of the *Snrpn* locus (Faulk et al. 2015; Kochmanski et al. 2018; Soubry et al. 2017; Susiarjo et al. 2013). Given that the *Srnpn* locus is hypermethylated in both blood and liver with gestational Pb exposure, it is plausible that it may represent an important biomarker of early life Pb exposure. We recently reported that perinatal BPA exposure leads to reprogramming of DNA methylation and hydroxymethylation at 12 imprinted genes (Kochmanski et al. 2018). Interestingly, 9 of the imprinted loci altered by Pb exposure were among those reprogrammed by BPA exposure. These data suggest that a core set of imprinted genes may be vulnerable to epigenetic reprogramming by multiple environmental exposures. As both Pb and BPA exposure have been implicated in adverse metabolic health outcomes (Bansal et al. 2017; Nadal et al. 2017; Niessen et al. 2018), it is plausible that reprogramming of genomic imprinting may represent a common mechanism linking both toxicants to these outcomes. The mechanistic consequences of environment-mediated dysregulation of imprinted genes, and the implications this may have for human health, should be investigated in future studies.

### Selection of surrogate tissues for environmental epigenomics studies

When comparing the chromosomal locations of DMCs/DMRs between blood and liver, only a small number of sites directly overlapped between the two tissues. Among DMRs/DMCs that overlapped between blood and liver, we identified several in males that exhibited changes in methylation in the same direction with Pb exposure in both blood and liver. These regions mapped to *Grifin, Elmsan1, Amn, Tbc1d30, Plekhg3*. An additional region did not map to a gene. Although the precise roles for the identified genes in normal physiology and disease are not fully characterized, recent evidence suggests they may have critical roles in development and in several disease states. (Griswold et al. 2011; Huyghe et al. 2013; Kalantry et al. 2001; Luder et al. 2008; Piraino and Furney 2017; Prisco et al. 2014). Although few differentially methylated sites overlapped directly, when comparing the lists of DMC-associated genes between blood and liver, we observed many DMC-associated genes that overlapped between tissues. The CpGs and genes identified in this study may therefore represent candidates for analysis in human environmental epigenetics studies. Future studies are needed to validate the plausibility of these loci as potential biomarkers of Pb exposure, as well as to investigate the functional and health-related consequences of altered DNA methylation at these sites.

### Limitations

This study has several important limitations. First, we utilized ERRBS, a method that, while providing resolution at the level of individual CpGs, does not cover the entire genome. The use of whole-genome bisulfite sequencing techniques may yield additional insight into the effects of Pb exposure on the liver, and identify additional Pb-induced signatures that overlap between blood and liver. Second, ERRBS does not differentiate between 5mC and 5hmC. Thus, the differentially methylated loci identified in this study are likely a combination of both 5mC and 5hmC. Third, changes of interest in DNA methylation may only occur in specific subpopulations of cells (Hui et al. 2018). Thus, our approach using unfractionated tissues may have missed other potential markers of interest for Pb exposure. Additionally, blood cells undergo rapid turnover, as well as exposure-, age-, and disease-dependent changes in cellular composition (Babio et al. 2013; Jaffe and Irizarry 2014; Su et al. 2016). Future studies using single-cell transcriptomics and epigenomics approaches will help to address issues of tissue-specific cellular heterogeneity and composition (Hui et al. 2018; Kelsey et al. 2017). Finally, it is important to note that DAVID gene ontology analysis does not consider the differing number of CpGs per gene, and therefore may exhibit bias.

### Conclusions

In summary, we have investigated the effects of perinatal Pb exposure on DNA methylation in paired samples of mouse blood and liver. There are several key strengths of this study, including the side-by-side comparison of blood and liver to identify signatures of Pb exposure present in the blood that reflect changes in the liver. Future studies should investigate the reproducibility of these findings, the mechanistic and functional consequences of persistent, Pb-induced changes in hepatic methylation, and the implications these changes have for human health.

## Acknowledgements

This work was supported by the National Institute of Environmental Health Sciences (NIEHS) TaRGET II Consortium (ES026697), along with the University of Michigan NIEHS/EPA Children’s Environmental Health and Disease Prevention Center P01 ES022844/RD83543601, the Michigan Lifestage Environmental Exposures and Disease (M-LEEaD) NIEHS Core Center (P30 ES017885), Institutional Training Grant T32 ES007062, and NIEHS grant R01 ES028802.

## Declaration

The authors declare no conflict of interest.

---

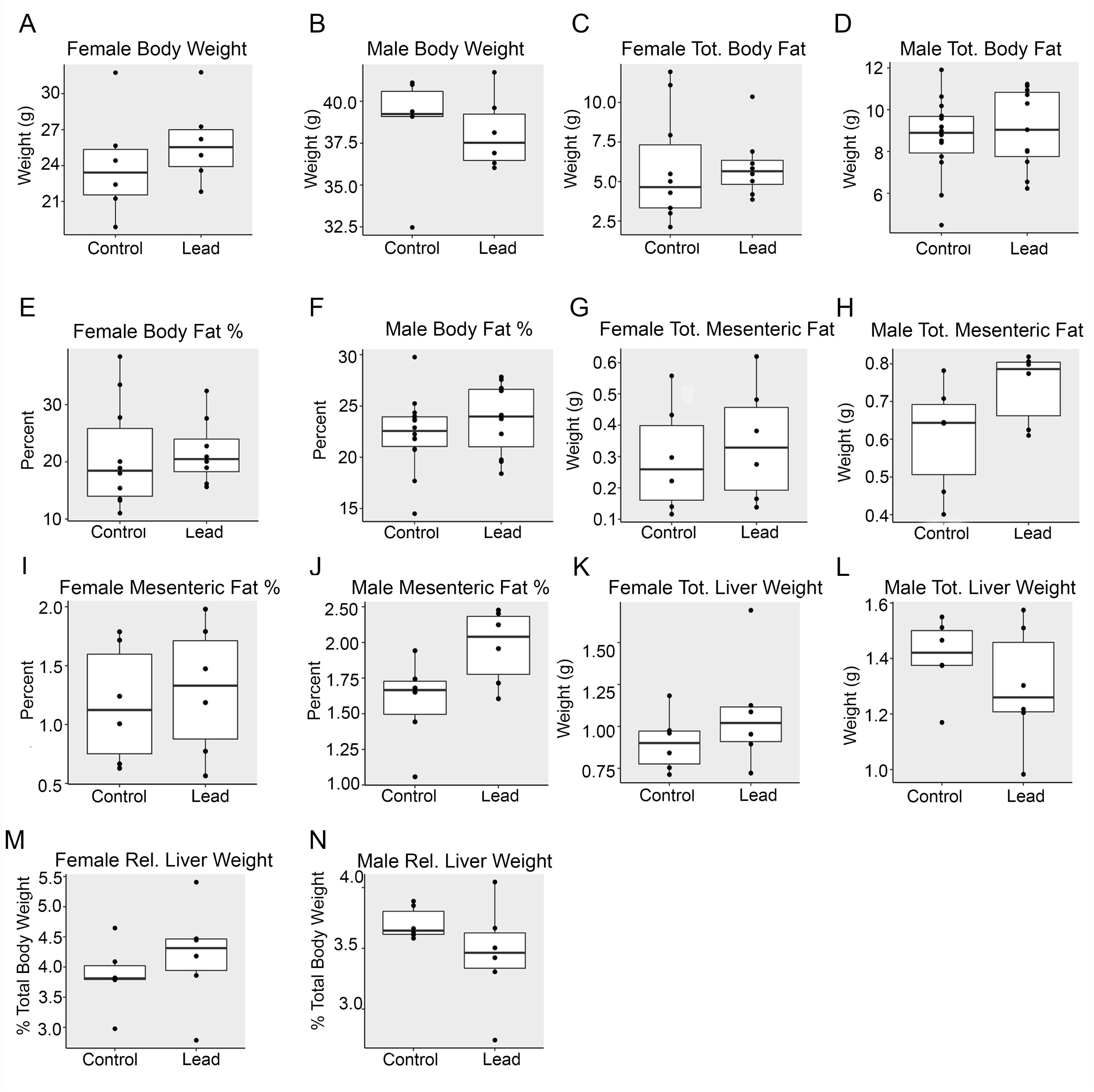

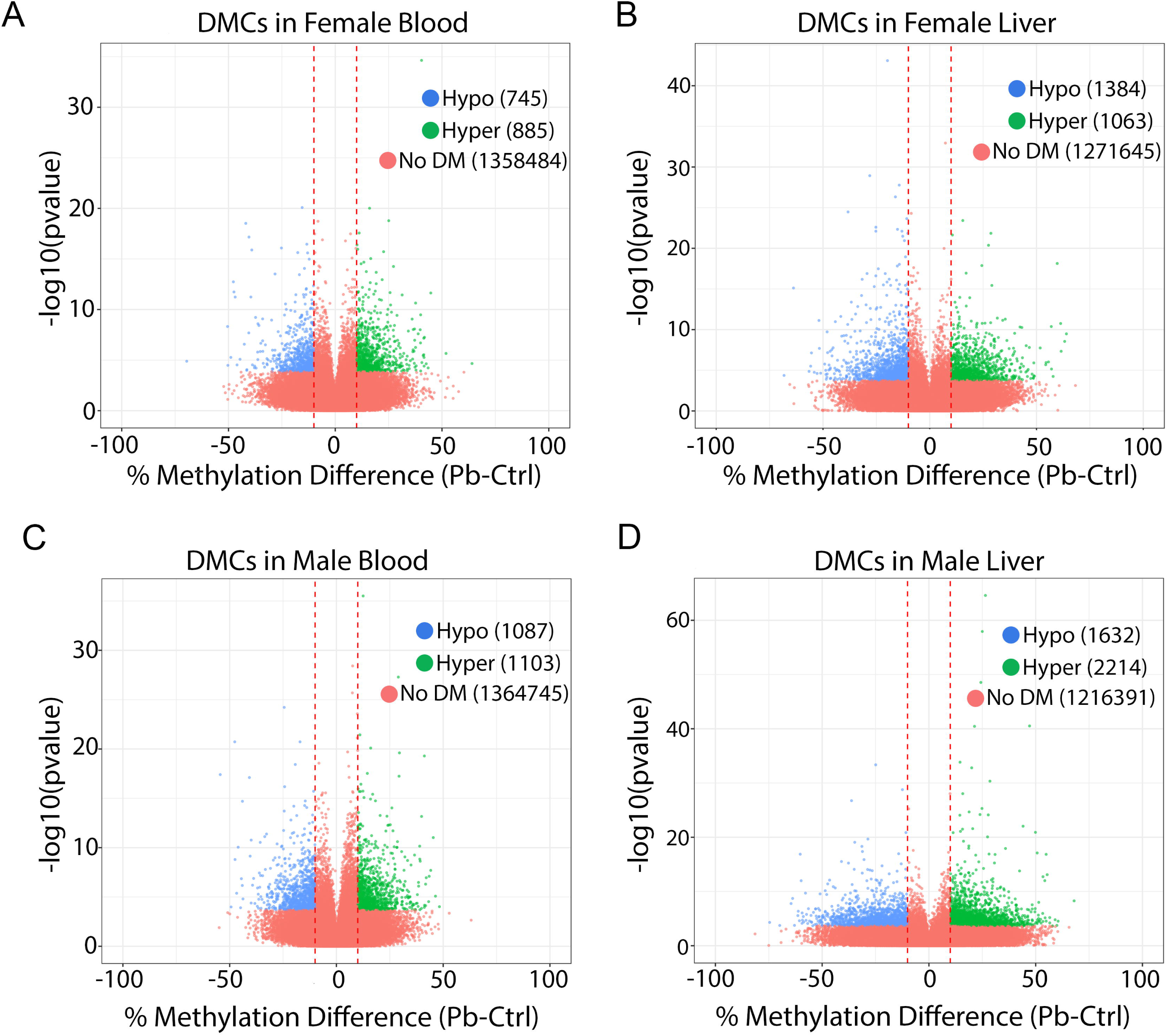

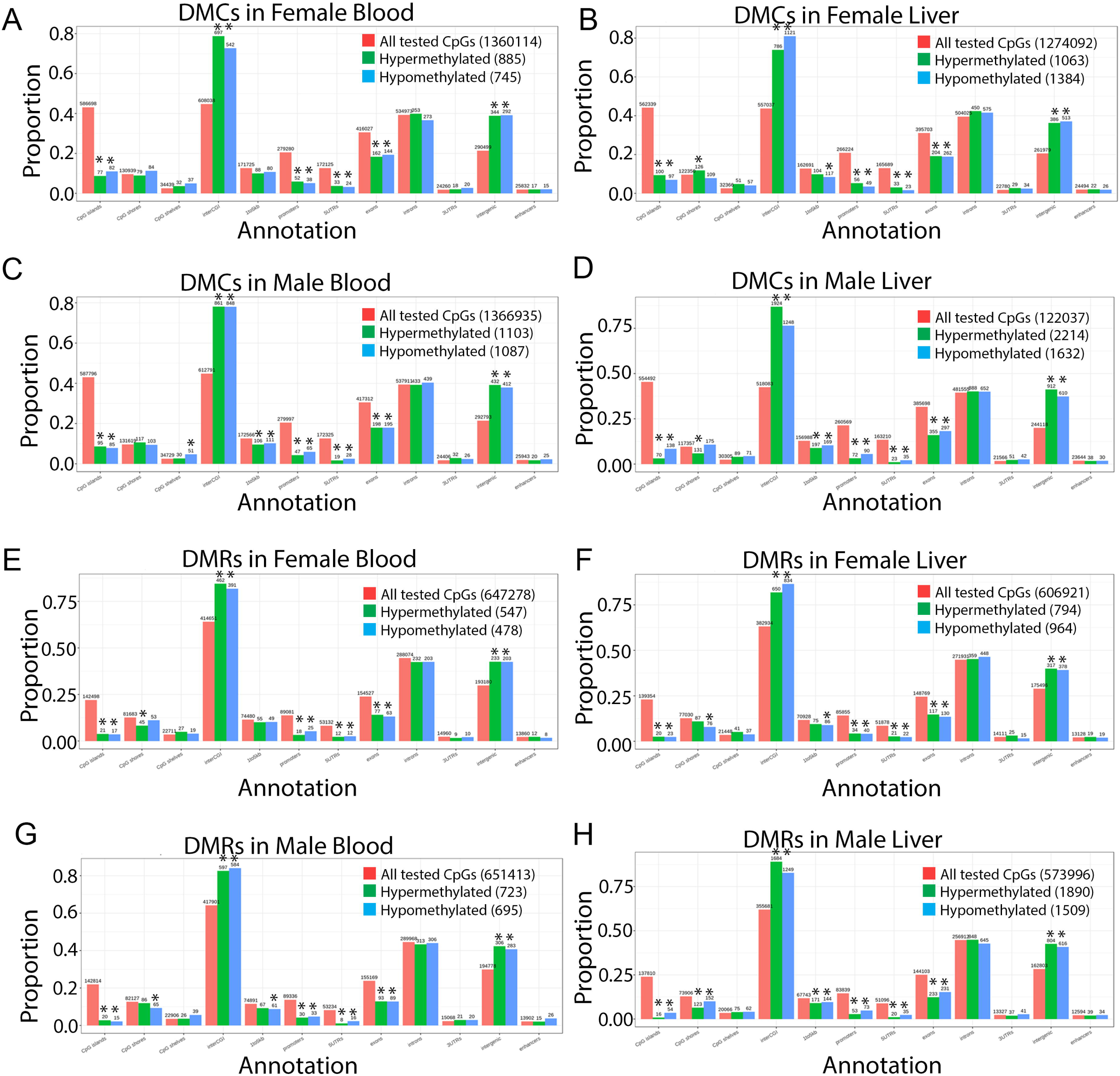

