## Supplemental Tables 1-8 for "Perinatal exposure to lead results in altered DNA methylation in adult mouse liver and blood: Implications for target versus surrogate tissue use in environmental epigenetics"

**Table S1**: Pathway analysis of differentially methylated regions identified in female blood

| Differentially hypomethylated promoter regions | | |
| --- | --- | --- |
| Pathway | FDR | Genes |
| GO:0060644 Mammary gland epithelial cell differentiation | 0.0002 | *Hoxa5*  *Prlr* |
| GO:0004896 Cytokine receptor activity | 0.0004 | *Il13ra1*  *Il3ra*  *Prlr* |
| GO:0061430 Bone trabecula morphogenesis | 0.050 | *Chad*  *Plxnb1* |
| Differentially hypermethylated promoter regions | | |
| Pathway | FDR | Genes |
| GO:0001191 Transcriptional repressor activity, RNA polymerase II transcription factor binding | 0.010 | *Cited1*  *Gbx2*  *Pou5f1*  *Dnajb1* |

Note: Pathway analysis conducted using Poly-Enrich (see Methods). An FDR<0.05 was considered significant.

**Table S2**: Pathway analysis of differentially methylated regions identified in female liver

| Differentially hypomethylated promoter regions | | |
| --- | --- | --- |
| Pathway | FDR | Genes |
| GO:1990204 Oxidoreductase complex | 0.009 | *Pdha1*  *Ndufa1*  *Noxa1* |
| Differentially hypermethylated promoter regions | | |
| Pathway | FDR | Genes |
| GO:0005326 Neurotransmitter transporter activity | 0.001 | *Slc18a3*  *Slc6a8* |

Note: Pathway analysis conducted using Poly-Enrich (see Methods). An FDR<0.05 was considered significant.

**Table S3**: Pathway analysis of differentially methylated regions identified in male blood

| Differentially hypomethylated promoter regions | | |
| --- | --- | --- |
| Pathway | FDR | Genes |
| GO:0015269 Calcium-activated potassium channel activity | 0.001 | *Kcnn1*  *Kcnk18* |
| GO:0009072 Aromatic amino acid family metabolic process | 0.003 | *Ido1*  *Iyd* |
| Differentially hypermethylated promoter regions | | |
| Pathway | FDR | Genes |
| None |  |  |

Note: Pathway analysis conducted using Poly-Enrich (see Methods). An FDR<0.05 was considered significant.

**Table S4**: Pathway analysis of differentially methylated regions identified in male liver

| Differentially hypomethylated promoter regions | | |
| --- | --- | --- |
| Pathway | FDR | Genes |
| GO:0090103 Cochlea morphogenesis | 3.16E-7 | *Gli2, Hpn, Wnt5a, Ptk7* |
| GO:0060026 Convergent extension | 4.59E-5 | *Wnt5a, Ptk7, Nkd1* |
| GO:0098810 Neurotransmitter reuptake | 6.08E-5 | *Slc6a3, Nos1, Rab3b* |
| GO:0042472 Inner ear morphogenesis | 6.30E-5 | *Dlx5, Gli2, Hpn, Otx1, Pou4f3, Wnt5a, Ptk7* |
| GO:0060071 Wnt signaling pathway, planar cell polarity pathway | 0.0001 | *Ryk, Wnt5a, Ptk7, Nkd1* |
| GO:0090162 Establishment of epithelial cell polarity | 0.0001 | *Wnt5a, Ptk7, Frmd4b, Myo18a* |
| GO:0030903 Notochord development | 0.0002 | *Gli2, Wnt5a, Noto* |
| GO:0032088 Negative regulation of NF-kappaB transcription factor activity | 0.0003 | *Cmklr1, Cmmd1, Rbck1, Dab2ip, Arrb2* |
| GO:0035066 Positive regulation of histone acetylation | 0.002 | *Lif, Nos1, Wbp2* |
| GO:1901021 Positive regulation of calcium ion transmembrane transporter activity | 0.003 | *Cacnb2, Kcne3, Plcg2* |
| GO:0050961 Detection of temperature stimulus involved in sensory perception | 0.006 | *Asic3, Arbb2* |
| GO:0090178 Regulation of establishment of planar polarity involved in neural tube closure | 0.006 | *Wnt5a, Ptk7* |
| GO:0032225 Regulation of synaptic transmission, dopaminergic | 0.006 | *Rab3b, Arbb2* |
| Differentially hypermethylated promoter regions | | |
| Pathway | FDR | Genes |
| GO:1902590 Multi-organism organelle organization | 0.0003 | *Chmp2a, Chmp6* |
| GO:0090185 Negative regulation of kidney development | 0.001 | *Ctnnb1, Pax8* |
| GO:0090178 Regulation of establishment of planar polarity involved in neural tube closure | 0.002 | *Celsr1, Dvl1* |
| GO:0001702 Gastrulation with mouth forming second | 0.002 | *Ctnnb1, Celsr1, Tenm4* |
| GO:0030825 Positive regulation of cGMP metabolic process | 0.002 | *Nos3, Rundc3a* |
| GO:0022010 Central nervous system myelination | 0.002 | *Kcnj10, Tenm4* |
| GO:0055062 Phosphate ion homeostasis | 0.002 | *Slc34a1, Gcm2* |
| GO:0015114 Phosphate ion transmembrane transporter activity | 0.003 | *Slc34a1, Slc17a7* |
| GO:0055075 Potassium ion homeostasis | 0.003 | *Kcnj10, Kcnma1* |
| GO:0006836 neurotransmitter transport | 0.003 | *Celsr1, Dvl1, Kcnj10, Slc17a7, Tc2n, Cplx4* |
| GO:0036452 ESCRT complex | 0.004 | *Chmp2a, Chmp6* |

Note: Pathway analysis conducted using Poly-Enrich (see Methods). An FDR<0.05 was considered significant.

**Table S5:** DMRs (differentially methylated regions) mapping to imprinted genes in female blood

| Gene | Start | End | Location | Change in methylation | FDR |
| --- | --- | --- | --- | --- | --- |
| *Meg3* | 109547801 | 109547850 | 1 to 5 kb | 36.92 | 0.002 |
| *Slc22a3* | 12454851 | 12454900 | Intron | -14.48 | 5.41E-05 |
| *Tnfrsf23* | 143670801 | 143670850 | Exon | -10.12 | 0.001 |

**Table S6:** DMRs (differentially methylated regions) mapping to imprinted genes in female liver

| Gene | Start | End | Location | Change in methylation | FDR |
| --- | --- | --- | --- | --- | --- |
| *Art5* | 102100151 | 102100200 | Promoter | 15.85 | 0.015 |
| *Blcap/Nnat* | 157561401 | 157561450 | 1 to 5 kb | -12.29 | 0.001 |
| *Fthl17e* | 9028751 | 9028800 | 1 to 5 kb | -14.42 | 0.0007 |
| *Gnas* | 174285401 | 174285450 | Exon/3’ UTR | 10.95 | 0.007 |
| *Grb10* | 12036901 | 12036950 | Intron | -11.08 | 7.05E-05 |
| *H19* | 142578751 | 142578800 | Promoter | -18.38 | 0.043 |
| *H19* | 142578951 | 142579000 | 1 to 5 kb | -23.47 | 0.025 |
| *Jade1* | 143395301 | 143395350 | Intron | 14.26 | 0.0003 |
| *Kcnq1* | 143395301 | 143395350 | Intron | 22.05 | 0.0002 |
| *Kcnq1* | 143295401 | 143295450 | Intron | -13.74 | 0.002 |
| *Kcnq1ot1* | 143295401 | 143295450 | Exon | -13.74 | 0.002 |
| *Magel2* | 62378951 | 62379000 | Exon | -20.11 | 0.017 |
| *Nespas* | 174285401 | 174285450 | Intron | 10.94 | 0.007 |
| *Pde10a* | 8865401 | 8865450 | Intron | -21.85 | 0.002 |
| *Pde10a* | 8754051 | 8754100 | Intron | -14.17 | 0.002 |
| *Pde4d* | 109659151 | 109659200 | Intron | -18.56 | 0.029 |
| *Tnfrsf23* | 143678051 | 143678100 | Intron | -24.14 | 0.038 |

Note: Methylation changes are expressed as lead vs. control.

**Table S7:** DMRs (differentially methylated regions) mapping to imprinted genes in male blood

| DMRs in Blood | | | | | |
| --- | --- | --- | --- | --- | --- |
| Gene | Start | End | Location | Change in methylation | FDR |
| *Airn* | 12822701 | 12822750 | Intron | -11.17 | 0.0005 |
| *Airn* | 12790851 | 12790900 | Intron | -24.69 | 0.043 |
| *Ampd3* | 110795651 | 110795700 | Intron/Exon | -11.12 | 0.040 |
| *Art5* | 102104301 | 102104350 | 1 to 5 kb | 23.58 | 0.022 |
| *Begain* | 109056051 | 109056100 | 1 to 5 kb | 10.21 | 0.033 |
| *Cdkn1c* | 143461101 | 143461150 | Promoter | 18.86 | 0.018 |
| *Commd1* | 22973901 | 22973950 | 1 to 5 kb/Intron/Exon | 18.67 | 0.006 |
| *Fbxo40* | 36990101 | 36990150 | Exon/Intron/5’UTR | 11.29 | 0.003 |
| *Grb10* | 12038651 | 12038700 | 1 to 5 kb | 11.31 | 0.019 |
| *Igf2r* | 12726401 | 12726450 | Intron | 17.14 | 0.006 |
| *Kcnq1* | 143377001 | 143377050 | Intron | 16.35 | 0.030 |
| *Peg12* | 62463901 | 62463950 | Exon | -23.40 | 0.003 |
| *Rasgrf1* | 89958351 | 89958400 | Intron | 19.19 | 2.62E-05 |
| *Scin* | 40133701 | 40133750 | Intron | 10.02 | 0.039 |
| *Snrpn* | 60012751 | 60012800 | Intron | 21.67 | 0.0001 |
| *Usp29* | 6770451 | 6770500 | Intron | -12.12 | 0.002 |
| *Zim1* | 6678051 | 6678100 | Exon | -12.05 | 0.0003 |
| *Zrsr1* | 22973901 | 22973950 | Exon | 18.67 | 0.006 |

Note: Methylation changes are expressed as lead vs. control.

**Table S8:** DMRs (differentially methylated regions)mapping to imprinted genes in male liver

| Gene | Start | End | Location | Change in methylation | FDR |
| --- | --- | --- | --- | --- | --- |
| *Cdkn1c* | 143459451 | 143459500 | Promoter/Intron | -23.11 | 0.003 |
| *Cdkn1c* | 143460301 | 143460350 | Promoter/Exon/1 to 5 kb | -11.38 | 0.024 |
| *Commd1* | 22978301 | 22978350 | Intron/Exon | -36.13 | 0.0006 |
| *Dlk1* | 109460001 | 109460050 | Intron/Exon | 12.84 | 0.048 |
| *Dlx5* | 6881401 | 6881450 | Intron | -16.01 | 0.012 |
| *H13* | 152696101 | 152696150 | Intron/Exon | -11.77 | 0.034 |
| *Htr2a* | 74706651 | 74706700 | Exon/3’UTR | 17.33 | 0.019 |
| *Kcnk9* | 72548401 | 72548450 | 1 to 5 kb | 16.96 | 0.048 |
| *Pde10a* | 8724551 | 8724600 | Intron | 30.84 | 0.004 |
| *Pde4d* | 108848451 | 108848500 | Intron | 19.29 | 0.022 |
| *Rasgrf1* | 89954101 | 89954150 | Intron/Exon | 10.15 | 0.017 |
| *Sfmbt2* | 10368301 | 10368350 | 1 to 5 kb | -28.39 | 0.003 |

**Figure S1:** Boxplots depicting phenotypic data from 5-month-old control and Pb-treated mice. Statistical significance was assessed using linear mixed effects regression. Body fat/mesenteric fat percentages and relative liver weights are expressed relative to total body weight.

**Figure S2:** Volcano plots showing differentially methylated cytosines (DMCs) for lead vs. control in female blood (A), female liver (B), male blood (C), and male liver (D). Green: regions significantly hypermethylated with Pb exposure. Blue: regions significantly hypomethylated with Pb exposure.

**Figure S3:** Summary plots depicting the total number of tested CpGs (pink), hypermethylated CpGs (green), and hypomethylated CpGs (blue) for each genomic annotation using the R annotatr package. Panels A-D depict differentially methylated cytosines (DMCs), and panels E-H depict differentially methylated regions (DMRs).
